# EMT activates ER-to-Golgi trafficking through upregulation of REEP2 to promote lung cancer progression

**DOI:** 10.1101/2025.11.12.688098

**Authors:** Kevin Fulp, Oluwafunminiyi Obaleye, Shike Wang, Xin Liu, Jiang Yu, Jonathan M. Kurie, Jun Xu, William K. Russell, Xiaochao Tan, Guan-Yu Xiao

## Abstract

Membrane trafficking governs the transport of molecules to both intracellular and extracellular locations, thereby maintaining cell homeostasis. During cancer progression, alterations in membrane trafficking are frequently observed. However, the mechanisms underlying the dysregulation of membrane trafficking in cancer progression remain largely unresolved. Recent evidence has demonstrated that epithelial-to-mesenchymal transition (EMT) in lung adenocarcinoma (LUAD) employs a membrane trafficking program to coordinate cancer cell invasion and immunosuppression in the tumor microenvironment (TME). To further dissect the pro-tumorigenic membrane trafficking program, here we conducted a CRISPR interference (CRISPRi) *in vivo* screen for membrane trafficking regulators in a syngeneic mouse model with a complete immune system. This screen identified REEP2, an endoplasmic reticulum (ER) shaping protein, as a novel regulator of the EMT-dependent membrane trafficking program, which is associated with a poor prognosis in LUAD patients and is required for LUAD metastasis in a syngeneic orthotopic LUAD mouse model. Mechanistically, the EMT activator ZEB1 upregulates REEP2 expression through miR-183- and miR-193a-mediated regulation that promotes the transportation of secretory cargoes from the ER exit site (ERES) to the Golgi, thereby augmenting the secretion of pro-tumorigenic factors. The REEP2-driven secretion promotes cancer cell proliferation, migration, and the infiltration of myeloid-derived suppressor cells (MDSCs) in the TME. These findings identify REEP2 as a critical mediator of the EMT-driven pro-metastatic membrane trafficking program, revealing a specific vulnerability in mesenchymal LUAD.

## Introduction

Membrane trafficking, a highly regulated process that is essential for cellular homeostasis, is disrupted on multiple levels in cancer, framing a new frontier in targeted therapeutics for cancer treatments ^1–7^. It involves the complex coordination of a host of vesicles shuttling an array of cargo to and from the endoplasmic reticulum (ER), Golgi apparatus, the plasma membrane, and extracellular space ^5–8^. Numerous cancer-associated mutations have been identified in the components of the membrane trafficking machinery ^9,10^. Furthermore, cancer cells coordinate membrane trafficking components post-transcriptionally (i.e. microRNA regulation) ^11–14^ and post-translationally (i.e. phosphorylation) ^15–20^ to dynamically adapt membrane trafficking pathways and acquire metastatic capabilities. Although numerous genome-wide analyses have been performed to interrogate the roles of membrane trafficking components in cancer cells ^21,22^, many critical knowledge gaps remain regarding the mechanisms of their dysregulation in cancer progression, and this area continues to be fertile ground for the discovery of novel cancer treatments.

Recent studies reveal that epithelial-to-mesenchymal (EMT)-activating transcription factors coordinate membrane trafficking networks to establish a front-rear polarity axis that facilitates cancer cell motility ^14,23^. Importantly, the EMT-driven membrane trafficking program underlies a vulnerability of EMT-driven metastatic cancers ^11,12^, and the inhibition of EMT-activated membrane trafficking regulators reactivates anti-tumor immunity in the tumor microenvironment (TME) and reverses acquired resistance to immune checkpoint inhibitor (ICI) therapy in LUAD ^11^. However, no comprehensive studies have systematically examined how membrane trafficking machinery influences TME that is essential for cancer progression.

To identify the membrane trafficking regulators critical for tumor growth with immune selective pressure, here we conducted a CRISPR repression (CRISPRi) *in vivo* screen in a syngeneic mouse model. Using LUAD cell lines derived from humans and genetically engineered mice that develop metastatic LUAD, which are classified as epithelial or mesenchymal ^11,24–26^ (Table S1), we further investigated regulatory mechanisms of the identified candidate, REEP2, an endoplasmic reticulum (ER) shaping protein, in membrane trafficking and LUAD progression. The findings presented herein uncover a molecular underpinning by which EMT promotes the secretory cargo transportation from the ER exit site (ERES) to the Golgi and, in turn, enhances the secretion of pro-tumorigenic factors that foster cancer cell motility and immunosuppressive TME.

## Results

### Pro-tumorigenic membrane trafficking program for LUAD growth

To systematically characterize the EMT-driven membrane trafficking in LUAD progression and immunosuppression, we transduced a CRISPR interference (CRISPRi) library targeting 2,099 trafficking-related genes ^27^ into murine mesenchymal LUAD 344SQ cells, which develop into metastatic LUADs owing to high expression of the EMT activator, ZEB1 ^26^ (Fig. 1A). We performed an *in vivo* screen by transplanting the cells into syngeneic wild-type (WT) immunocompetent mouse flanks and measured the effects on tumorigenesis (Fig. 1A). This screen revealed a panel of membrane trafficking regulators required for tumor growth under immune surveillance (fold-change (log_2_) < -1, FDR (-log_10_) > 20) (Fig. 1B, blue box, Table S2). We prioritized a set of top hits (fold-change (log_2_) < -2 and FDR (-log_10_) > 20, Fig. 1B, red dots and Fig. 1C) and assessed their correlation with EMT (Fig. 1D) and prognostic values (Fig. 1E and Fig. S1) using The Cancer Genome Atlas (TCGA) database. We identified several membrane trafficking regulators (*KRT16, REEP2, TNNT1, ARL14*) that significantly correlate with the EMT signature and shorter survival durations of LUAD patients. These findings reveal a membrane trafficking program that contributes to LUAD tumorigenesis. Among these hits, REEP2 represents the strongest EMT correlation and demonstrates high prognostic value, and correlates with LUAD metastasis (Fig. 1F).

**Figure 1.**
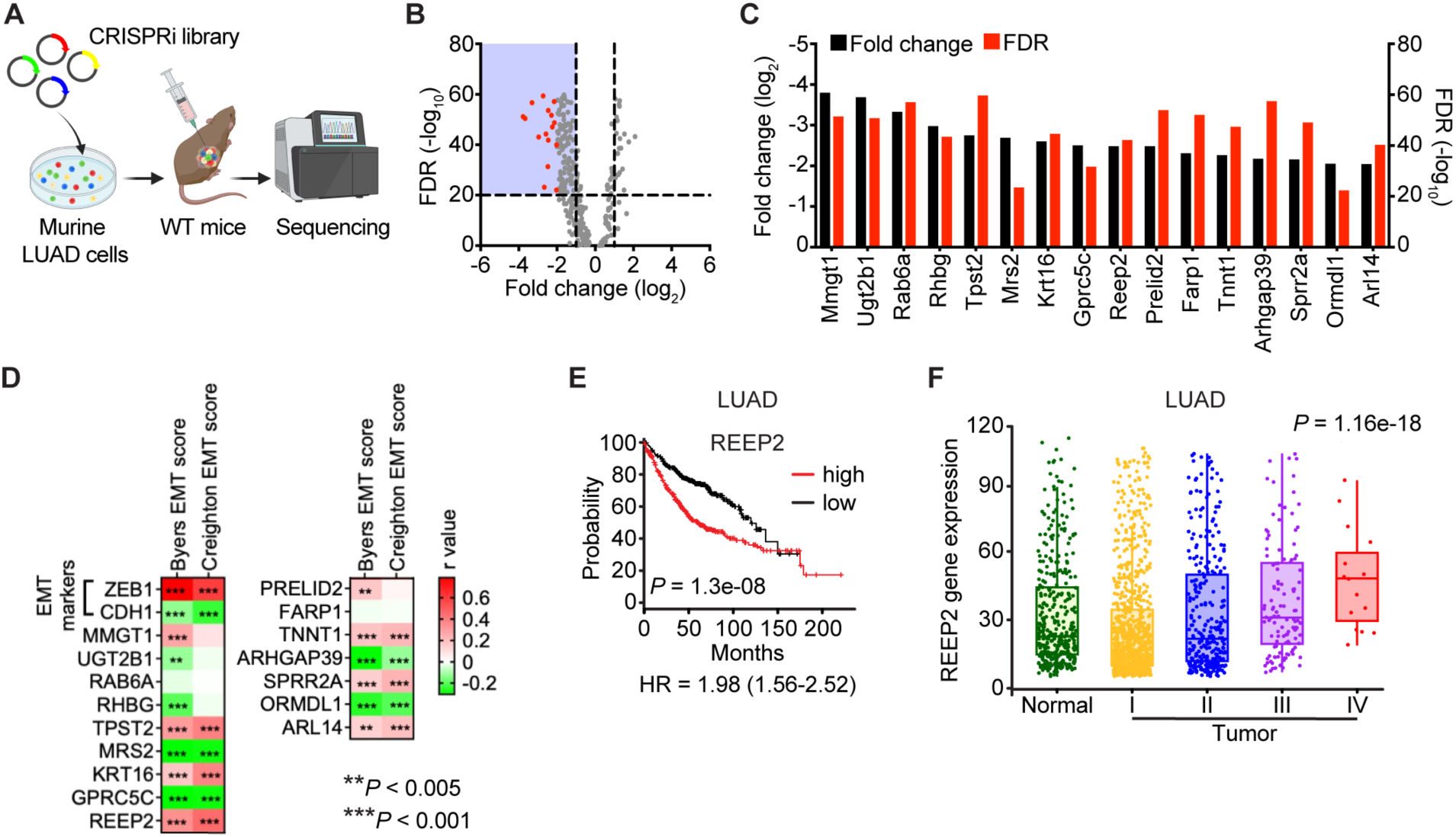
Pro-tumorigenic membrane trafficking program for LUAD growth. (A) A CRISPRi library targeting membrane trafficking genes was transduced into 344SQ murine LUAD cells. Cells were transplanted subcutaneously on the flanks of WT syngeneic mice. Following tumor formation, the tumors were harvested and sgRNAs were sequenced. (B) Volcano plots of the analyses of the screening results in immunocompetent WT mice. Grey dots represent membrane trafficking genes, and vertical and horizontal dotted lines indicate fold-change (log_2_) = ±1 and FDR (-log_10_) = 20, respectively. The red dots represent the top hits that fold-change (log_2_) < -2 and FDR (-log_10_) > 20. (C) The bar graph represents the top hits in (B) (Red dots). (D) Heat map illustration of correlation between mRNA levels of indicated genes and EMT scores (Byers or Creighton) in the TCGA LUAD cohort. r values: Pearson correlation. (E) Kaplan-Meier survival analysis of LUAD patients based on REEP2 mRNA levels above (high) or below (low) the median value. (F) Expression levels of REEP2 mRNA in normal lung tissues (n = 391), stage I (n = 905), stage II (n = 331), stage III (n = 146), and stage IV lung tumors (n = 17) using data from TNMplot. (https://tnmplot.com/).

### REEP2 is required for mesenchymal LUAD metastasis

To further investigate the function of REEP2 in LUAD progression, we subjected murine mesenchymal 344SQ cells to CRISPR/Cas9-mediated REEP2 knockout (REEP2 KO) (Fig. S2A), and inoculated the cells orthotopically in the lungs of the syngeneic WT mice. We found that REEP2 KO cells generated orthotopic LUADs that were smaller (Fig. 2A, left dot plot) and less metastatic to the mediastinum (Fig. 2A, middle dot plot) and the contralateral lung (Fig. 2A, right dot plot). We then assessed cancer progression properties *in vitro* and found that REEP2 depletion (Fig. S2A-D) reduced murine (344SQ) and human (H1299) mesenchymal LUAD cell proliferation (Fig. 2B), but negligibly affected the viability of human epithelial LUAD (HCC827) (Fig. S2E) or non-cancerous human bronchial epithelial cell (HBEC30KT) (Fig. S2F). Furthermore, we found that REEP2 deficiency in mesenchymal LUAD cells resulted in cell cycle arrest (Fig. 2C), with a negligible change in apoptosis (Fig. 2D), and sharply reduced migratory and invasive activities in Boyden chambers (Fig. 2E and F). These data indicate a significant, previously unknown association of REEP2 with LUAD progression.

**Figure 2.**
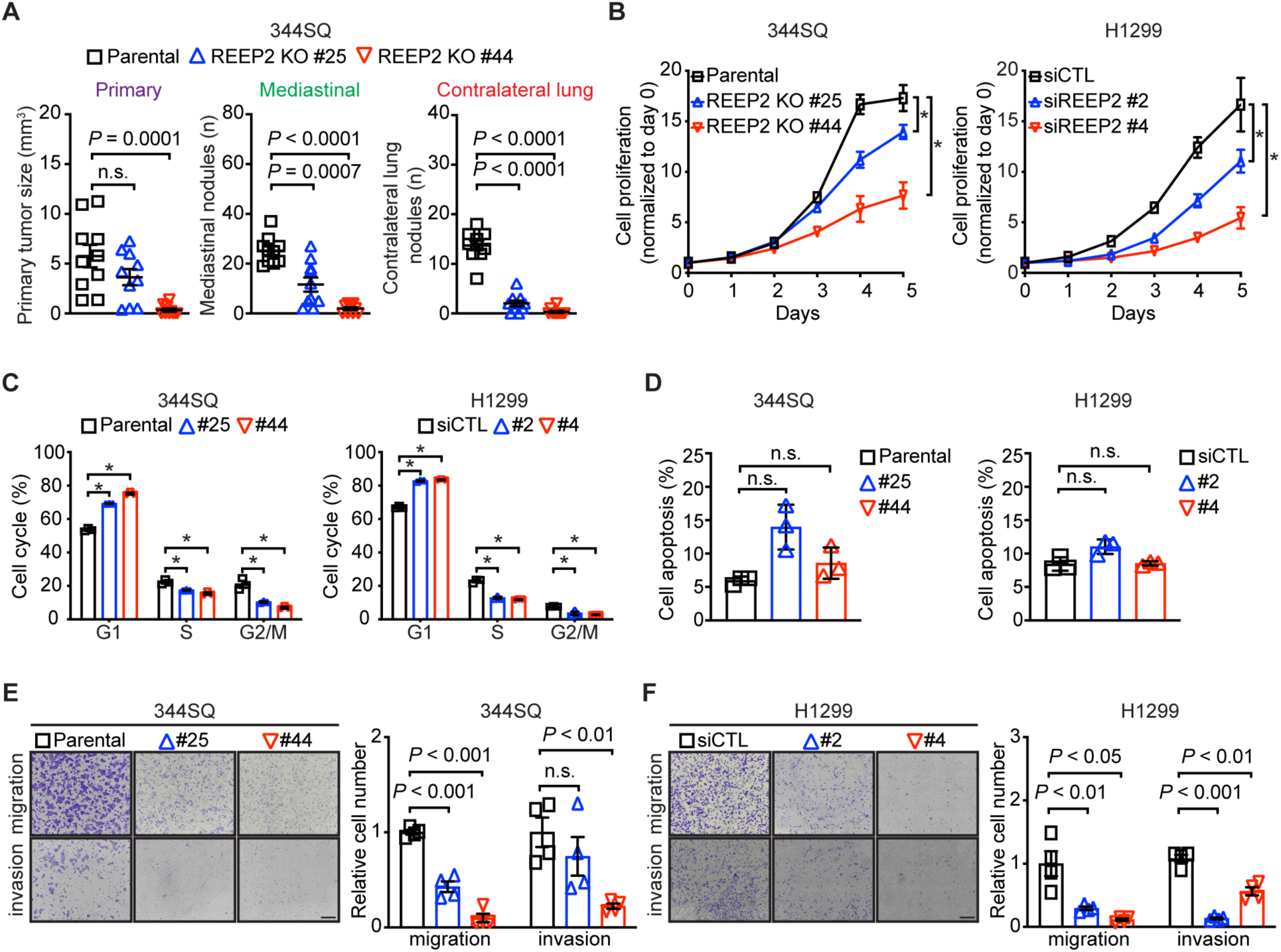
REEP2 is critical for EMT-driven LUAD progression. (A) Syngeneic, immunocompetent mice were injected with parental or REEP2 KO 344SQ cells and necropsies were performed 2 weeks later to assess tumor formation and metastasis (n = 10 mice per cohort). Orthotopic lung tumor size (left dot plot) and numbers of metastases to mediastinum (middle dot plot) and contralateral lung (right dot plot). (B) *In vitro* cell proliferation assay in murine 344SQ parental and REEP2 knockout (KO) cells, and human H1299 cells transfected with control siRNA (siCTL) or REEP2 siRNA (siREEP2). (C, D) Analyses of cell cycle (C) and apoptosis (D) in murine 344SQ parental and REEP2 knockout (KO) cells, and human H1299 cells with siRNA-mediated REEP2 knockdown (siREEP2). (E, F) Bright-field micrographs of crystal violet-stained murine (344SQ) (E) and human (H1299) (F) cells that migrated through noncoated (migration) or Matrigel-coated (invasion) filters in Boyden chambers. The scatter plots represent the quantification of replicates 16 h after seeding and normalized to parental. Results represent means ± SEM. *P* values were determined using two-tailed Student’s t-test.

### ZEB1 relieves REEP2 from microRNA-dependent silencing

We identified a positive correlation between ZEB1 and REEP2 expressions in human LUAD (Fig. 3A) and validated that their mRNA and protein levels changed in a corresponding fashion following ZEB1 gain- or loss-of-function (Fig. 3B, S3A). ZEB1 has been shown to silence microRNAs (miRs) to relieve the negative transcriptional regulation from genes involved in LUAD progression ^11,14,24,25^. Using the TargetScan prediction tool (www.targetscan.org), we identified predicted miR-183 and miR-193a binding sites on the REEP2 3’-untranslated regions (3’-UTRs) (Fig. S3B). In the epithelial cells with inducible ZEB1 expression (H322_iZEB1), ZEB1 induction repressed the expressions of both miR-183 and miR-193a, while upregulating REEP2 simultaneously (Fig. S3C). We found that miR-183 and miR-193a mimics reduce REEP2 expression in murine and human mesenchymal LUAD cells (Fig. 3C, S3D). To validate the miR-183- and miR-193a-mediated REEP2 silencing, we mutated the predicted binding sites (Fig. S3B) and found that both miR-183 and miR-193a mimics suppressed the activity of human REEP2 reporters containing wild-type but not mutant 3’-UTRs lacking predicted binding sites (Fig. 3D and E). These findings define a REEP2 regulatory axis in which ZEB1 alleviates miRNA-mediated repression, leading to REEP2 upregulation in mesenchymal LUAD.

**Figure 3.**
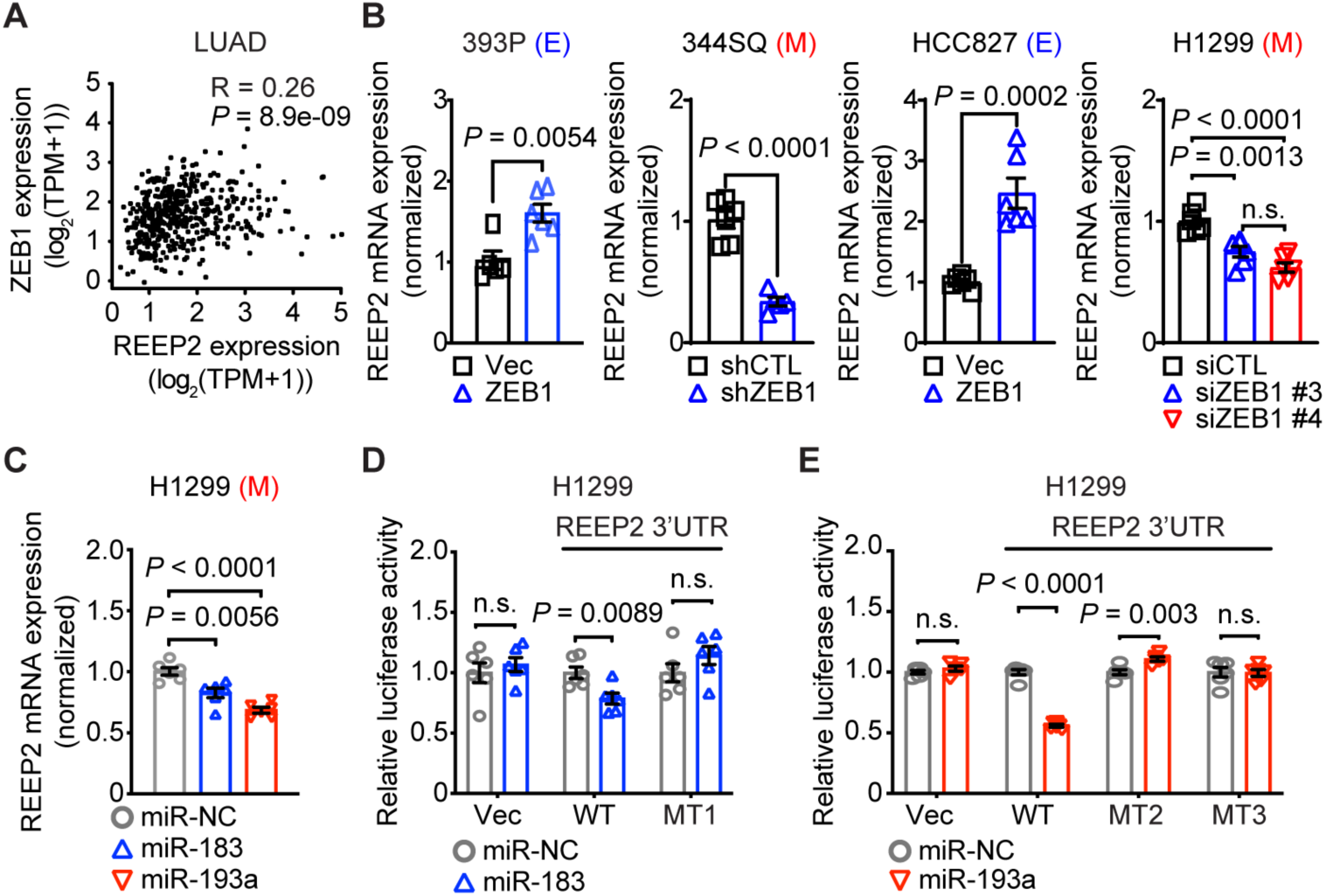
ZEB1 activates REEP2 via miRNAs. (A) The correlation between REEP2 and ZEB1 mRNA expression (TPM (transcripts per million)) in TCGA LUAD tissues. R, Spearman correlation. (B) qPCR analysis of REEP2 mRNA expression levels in murine (393P, 344SQ) and human (HCC827, H1299) LUAD cells with ectopic ZEB1-expression or ZEB1-depletion (n=6 replicates per condition). Vec: empty vector. E: epithelial. M: mesenchymal. (C) qPCR analysis of REEP2 mRNA expression levels in miR mimic-transfected human LUAD cells. NC: non-coding control. M: mesenchymal. (D, E) Human REEP2 3’-UTR reporter assays. Luciferase activities in H1299 cells transiently co-transfected with pre-miRs of miR-183 (D) or miR-193a (E) and human REEP2 3’UTR reporters containing wild-type (WT) or mutant 3’-UTRs lacking predicted miR-binding sites (MT1-3). Vec: empty vector. Results represent means ± SEM. *P* values were determined using two-tailed Student’s t-test.

### ZEB1 coordinates REEP2 to promote ERES and Golgi colocalization in mesenchymal LUAD

REEP proteins are known to generate membrane curvature of the ER to regulate its structure and function ^28^ and have been suggested to function in secretory trafficking ^29^. Proteins destined for secretion are synthesized in the ER, where cargoes are packaged into ERES and transferred to the Golgi apparatus ^30^, after which they are directed to the plasma membrane through post-Golgi traffic ^31^. SEC proteins, including SEC16 and SEC24, are recruited to the site of ERES and are essential for the formation of ERES and cargo packaging ^32–34^. GM130 at the cis-Golgi tethers ERES cargo delivery to the Golgi ^35^. Given the significant curvature at the ERES, we hypothesized that ZEB1 coordinates REEP2 to direct ERES toward the Golgi in mesenchymal LUAD (Fig. 4A). To test the possibility, we visualized the distribution of SEC16A and GM130 using immunofluorescence. Strikingly, compared to empty vector control (Vec), which showed the dispersed SEC16A signals outside of the Golgi area marked by GM130 (Fig. S4A, arrows), ectopic ZEB1 expression remarkably increased SEC16A colocalization with GM130 (Fig. S4A, arrowheads) in the epithelial murine (393P) LUAD cells. The depletion of ZEB1 in the mesenchymal murine (344SQ) and human (H1299) LUAD cells reduced the SEC16A/GM130 colocalization (Fig. S4B and C). The deficiency of REEP2 similarly decreased their colocalization in the mesenchymal human (H1299) LUAD cells (Fig. S4C). Furthermore, the ectopic expression of ZEB1 in the epithelial human (HCC827) LUAD cells increased the SEC16A/GM130 colocalization (Fig. 4B, Vec vs. siCTL), while REEP2 depletion (Fig. S4D) abolished the phenotype (Fig. 4B, siCTL vs. siREEP2s). We then observed the consistent findings of higher colocalization between SEC24D-labelled ERES and the Golgi in the epithelial human (HCC827) LUAD cells with ectopic ZEB1 expression compared to Vec (Fig. 4C), while REEP2 depletion disrupted the phenotype (Fig. 4D), suggesting ZEB1 coordinates REEP2 to promote ERES locating on the Golgi in mesenchymal LUAD.

**Figure 4.**
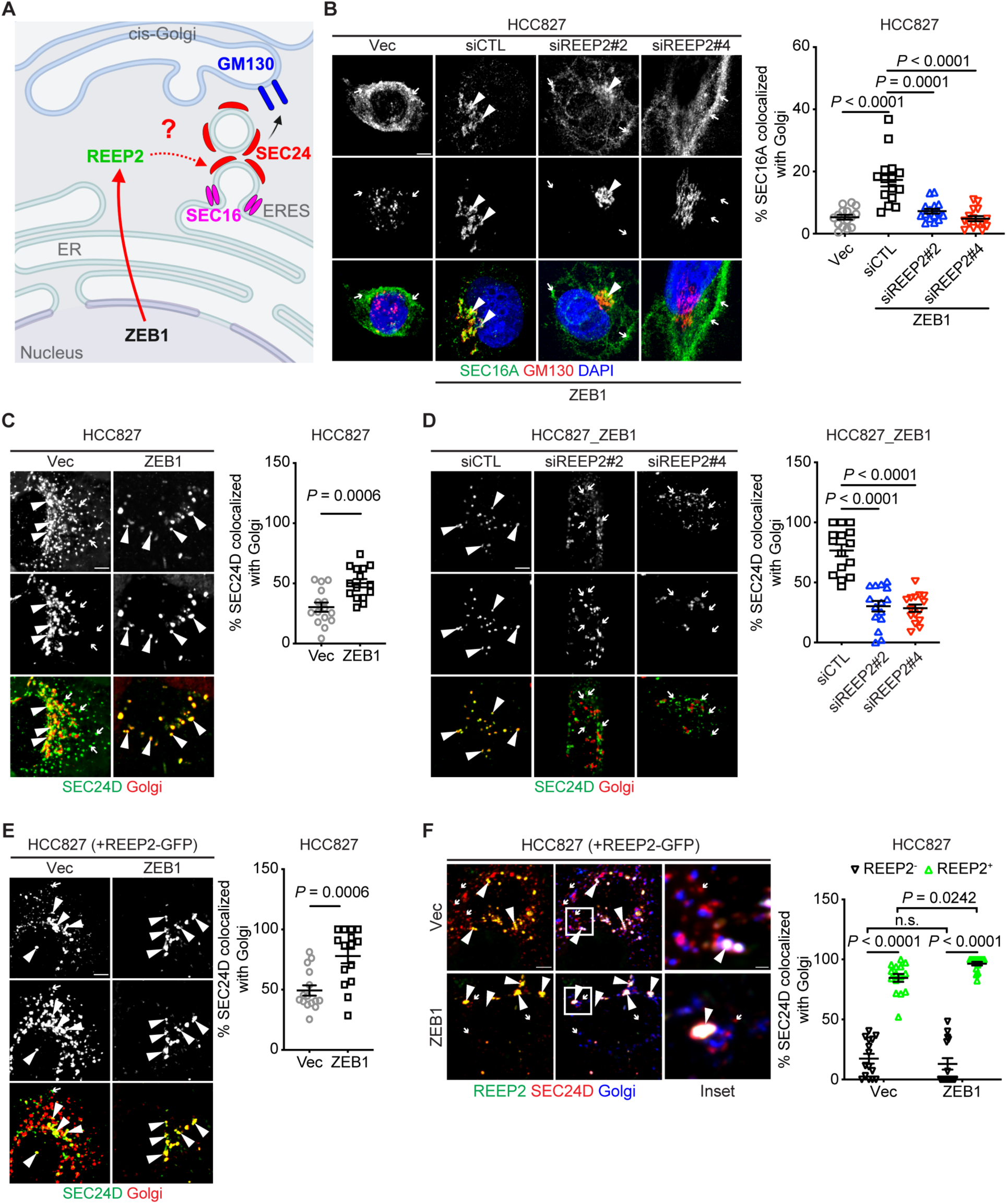
ZEB1 coordinates REEP2 to promote ERES colocalization with the Golgi. (A) Schematic illustration of a secretory pathway model. ZEB1 upregulates REEP2 to promote ERES (SEC16, SEC24) transportation to the Golgi (GM130). (B) Confocal micrographs of cells co-stained with anti-SEC16A (green) and anti-GM130 (red) antibodies. DAPI (blue). Scale bar: 5 μm. The scatter plots quantify Golgi-localized SEC16A per cell (dot) based on % of total SEC16A that co-localizes with Golgi (GM130 channel) (n = 15 cells per group). (C-E) Confocal micrographs of cells co-transfected with SEC24D-mCherry (green) and Golgi-pmTurquoise2 (red). DAPI (blue). Scale bar: 5 μm. The scatter plots quantify Golgi-localized SEC24D per cell (dot) based on % of total SEC24D that co-localizes with Golgi (n = 15 cells per group) in HCC827 cells with ectopic ZEB1-expression (C), and with REEP2 depletion (D) or REEP2-GFP transfection (E). (F) Confocal micrographs of cells co-transfected with REEP2-GFP (green), SEC24D-mCherry (red) and Golgi-pmTurquoise2 (blue). Scale bar: 5 μm, 1 μm (inset). The scatter plots quantify Golgi-localized SEC24D per cell (dot) based on % of total SEC24D that co-localizes with Golgi (n = 15 cells per group) in HCC827 cells with REEP2-GFP transfection. Results represent means ± SEM. *P* values were determined using two-tailed Student’s t-test.

Thus far, REEP2 was observed to form vesicular structures that lack colocalization with markers of canonical cellular compartments ^36^. To further investigate the distribution and function of REEP2, we transfected the HCC827 cells with fluorescent-tagged REEP2 (Fig. S4E) and found that the REEP2 vesicles highly colocalized with SEC24D-labelled ERES (∼80%) and there was no difference between ectopic Vec and ZEB1 expression (Fig. S4F). However, the ectopic expression of ZEB1 significantly enhanced the ERES colocalization with the Golgi (Fig. 4E). Strikingly, we observed that the SEC24D structures with REEP2 coating dramatically increased their colocalization with the Golgi, compared to those lacking REEP2 coating, and ectopic ZEB1 expression further raised the colocalization (Fig. 4F). These findings suggest that REEP2 recruitment to the ERES is necessary for their transportation to the Golgi.

### REEP2-driven ERES/Golgi colocalization promotes secretory trafficking

To test how REEP2-driven ERES/Golgi colocalization influences protein trafficking, we used the retention using selective hooks (RUSH) system ^37^ to quantitatively monitor trafficking kinetics at different time points. Human mesenchymal LUAD cells (H1299) were transfected with a GFP-tagged CD-MPR, a canonical cargo that localizes to the trans-Golgi network (TGN) ^38^, which binds to streptavidin fusing with an ER retention signal (Lys-Asp-Glu-Leu; KDEL) via the streptavidin-binding peptide (SBP) (Fig. S5A). Upon biotin treatment, the CD-MPR-RUSH is released from the ER and trafficked to the Golgi (Fig. 5A, siCTL, S5A). REEP2 depletion drastically reduced the trafficking of CD-MPR-RUSH to the Golgi at both 10 minutes and 30 minutes post-Biotin treatment (Fig. 5A).

**Figure 5.**
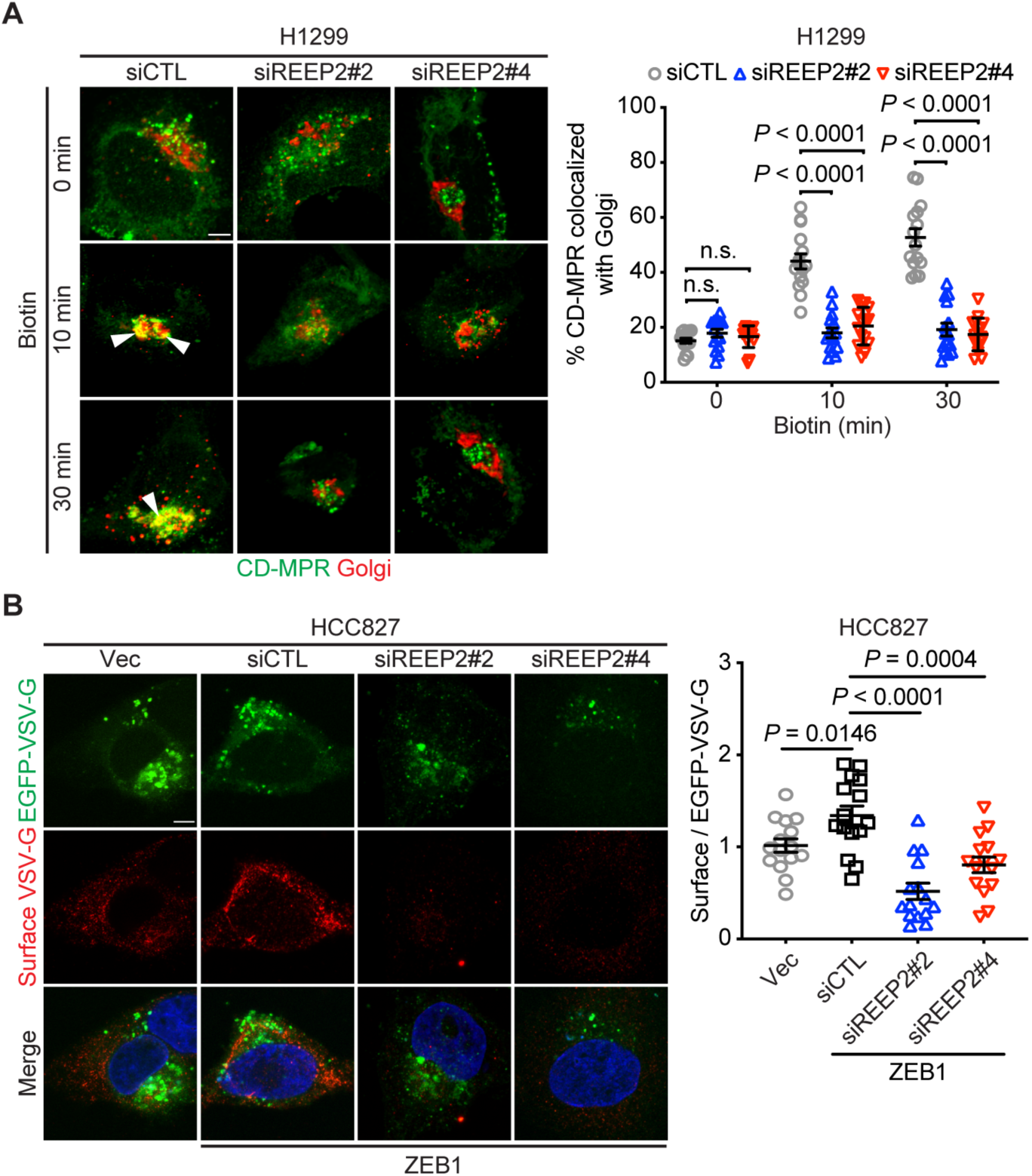
REEP2 promotes ER-to-Golgi trafficking. (A) Confocal micrographs of cells co-transfected with CD-MPR-RUSH-GFP (green) and Golgi-pmTurquoise2 (red). Scale bar: 5 μm. The scatter plots quantify Golgi-localized CD-MPR per cell (dot) based on % of total CD-MPR that co-localizes with Golgi at different time points after biotin treatment (40 μM) (n = 15 cells per group) in H1299 cells transfected with control siRNA (siCTL) or REEP2 siRNA (siREEP2). (B) Confocal micrographs of EGFP-VSV-G-transfected cells taken 1 h after transfer to permissive temperature. Scale bar, 5 μm. The scatter plot represents the ratio of surface VSV-G to EGFP-VSV-G in each cell (dot) (n = 15 cells per group) in HCC827 cells co-transfected with empty vector (Vec) or ZEB1 with control siRNA (siCTL) or REEP2 siRNA (siREEP2). Results represent means ± SEM. *P* values were determined using two-tailed Student’s t-test.

Next, to assess whether ERES/Golgi colocalization facilitates secretory trafficking, the ER stress inducer, thapsigargin, was used to disperse SEC16A ^34^, and the temperature-sensitive mutant vesicular stomatitis virus (VSV-G) assay was used to measure secretory trafficking (Fig. S5B). Consistent with the previous study ^34^, thapsigargin disrupted ERES colocalization with the Golgi (Fig. S5C). We observed that the reduced ERES/Golgi colocalization impacted secretory trafficking in the mesenchymal human (H1299) LUAD cells (Fig. S5D). We found that ZEB1 promoted secretory trafficking in the epithelial human (HCC827) LUAD cells and that the depletion of REEP2 abated the phenotype (Fig. 5B). These findings suggest that REEP2 coordinates ERES/Golgi colocalization to promote secretory trafficking.

### REEP2 regulates the secretion of pro-tumorigenic factors

Given that REEP2 promotes secretory trafficking, we hypothesized that REEP2 facilitates the secretion of pro-metastatic factors in mesenchymal LUAD cells. To test this, we evaluated whether REEP2-dependent secretions could modulate epithelial LUAD cell behaviors in a paracrine manner. Conditioned medium (CM) from REEP2-replete HCC827_ZEB1 cells induced a decrease in the G₀/G₁ population and a corresponding increase in S-phase of HCC827_Vec cells (Fig. 6A), and enhanced the cell mobility (Fig. 6B), whereas CM from REEP2-deficient HCC827_ZEB1 cells demonstrated no effects (Fig. 6A and B). These findings suggest that the REEP2-driven secretome contributes to the metastatic properties of LUAD cells.

**Figure 6.**
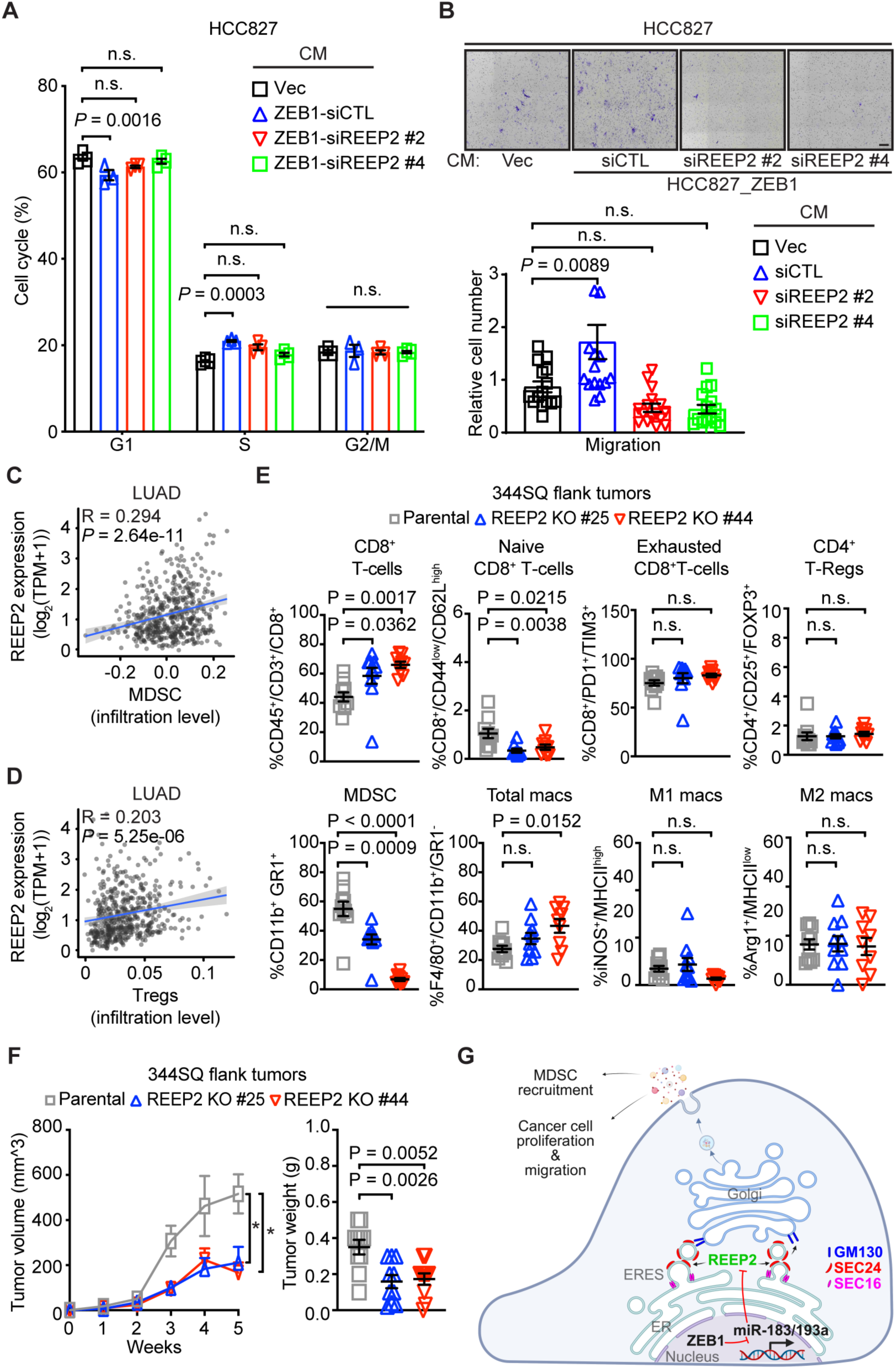
REEP2-driven secretome promotes LUAD progression. (A) Analyses of cell cycle in HCC827 cells treated with CM from indicated cells. (B) Bright-field micrographs of crystal violet-stained HCC827 cells that migrated through noncoated filters in Boyden chambers. Upper chambers loaded with indicated CM samples. The scatter plots represent the quantification of replicates 16 h after seeding and normalized to Vec. (C, D) Correlation of REEP2 mRNA levels with infiltration levels of myeloid-derived suppressor cells (MDSCs) (C) and regulatory T (Tregs) cells (D) quantified by quanTIseq analysis in a human LUAD cohort (Data generated from TIMER2.0). R, Spearman correlation. (E) Flow cytometric analysis of parental or REEP2-depleted (REEP2 KO #25 or #44) 344SQ-generated flank tumors isolated from syngeneic, immunocompetent mice (n = 10 tumors per cohort). Percentages of T cells (CD8^+^ T cells, CD44^low^/CD62L^high^ naïve CD8^+^ T cells, PD1^+^/TIM3^+^ exhausted CD8^+^ T cells and CD25^+^/FOXP3^+^ CD4^+^ Treg cells), MDSCs, and antigen-presenting cells (total macrophages, M1 macrophages, and M2 macrophages) were quantified. (F) Flank tumor volume (left) and weight (right) in syngeneic, immunocompetent mice (dots) injected with parental or REEP2-depleted (REEP2 KO #25 or #44) 344SQ cells. (G) Schematic illustration of a working model. ZEB1 relieves REEP2 from miR-183- and miR-193a-mediated silencing, thereby locating ERES on the Golgi and facilitating pro-metastatic secretion that recruits MDSC in the TME and promotes cancer progression.

To explore REEP2-dependent secreted factors, we performed liquid chromatography-mass spectrometry (LC-MS) analysis on CM collected from REEP2-replete HCC827_ZEB1 cells, and their REEP2-depleted counterparts. We found that REEP2 depletion reduced secretion of factors involved in the immunomodulatory and cell migratory pathways (Fig. S6A, blue and red dots) (Dataset S1), defined by enrichment analysis (Fisher’s exact test using Gene Ontology terms) of the down-regulated secreted proteins (Fig. S6B), including autotaxin (ATX). We have recently identified ATX as a predominant secretory factor in the EMT-driven secretion that promotes LUAD cell motility in autocrine and paracrine mechanisms and mediates the immunosuppressive TME^11^. Enzyme-linked immunosorbent assays (ELISA) on CM samples confirmed that REEP2 depletion reduces the secretion of ATX (Fig. S6C), whereas the intracellular ATX remains unchanged (Fig. S6D).

Given that we identified numerous immunomodulatory proteins that were diminished by REEP2 depletion (Fig. S6A, red dots), we speculate that REEP2-regulated secretome remodels TME. We used TIMER2.0 ^39–41^ to analyze the immune infiltrates in human LUAD cataloged in TCGA. We found that high REEP2 expression positively correlates with higher infiltration levels of myeloid-derived suppressor cells (MDSCs) and regulatory T cells (Tregs) in human LUAD (Fig. 6C and D). Therefore, we reasoned that REEP2 may create an immunosuppressive TME and addressed this possibility by carrying out flow cytometry on flank tumors generated by REEP2-replete or -deficient murine LUAD cells in syngeneic, immunocompetent mice. We found that REEP2 depletion enhanced total CD8^+^ T cells but reduced naïve CD8^+^ T cells and MDSCs (Fig. 6E) in the TME, and suppressed tumor growth (Fig. 6F). These findings suggest that REEP2 promotes the secretion of pro-tumorigenic factors required for LUAD progression.

## Discussion

Growing evidence shows that alterations in membrane trafficking frequently occur during cancer progression and, in turn, promote metastasis ^42^, indicating significant therapeutic potential for these pathways. Here, we show that EMT initiates a membrane trafficking program essential for LUAD tumorigenesis under immunosurveillance and identify that ZEB1 coordinates REEP2 via miR-183/miR-193a to enhance ERES colocalization with the Golgi, thereby facilitating the secretion of pro-metastatic factors that promote LUAD metastasis (Fig. 6G). This regulatory axis discloses a potential therapeutic vulnerability in EMT-driven metastatic cancers.

The REEP family comprises six ER-localized membrane-shaping proteins that facilitate high membrane curvature. REEP1-4 share structural motifs and are less abundant than REEP5 and 6, which are more broadly expressed ^28^. Germline mutations in REEP1 and REEP2 cause hereditary spastic paraplegia (HSP) and related motor neuropathies ^43,44^, underscoring their physiological importance in ER shaping and neuronal maintenance. Additionally, REEP family members have been implicated in tumor biology. REEP4 is upregulated in renal carcinoma, where its expression aligns with tumor aggressiveness ^45^, and REEP6 expression correlates with poor prognosis in oral squamous cell carcinoma ^46^. Moreover, REEP5 and REEP6 have been shown to potentiate CXCR1/IL-8 signaling and promote proliferation and metastasis in lung cancer cells^47^. These findings suggest that REEP family proteins broadly contribute to tumor progression, though their mechanisms and context-specific roles are still elusive. Here, we show for the first time that REEP2 is a downstream effector of ZEB1-dependent transcription program that activates ER-to-Golgi trafficking of secretory cargoes required for EMT-driven metastasis in LUAD.

A recent study demonstrated that REEP1-4 form a distinct ER-derived vesicular compartment and regulate ER tubule dynamics ^43,44^, but their roles in membrane trafficking remain largely unexplored. Here, we demonstrate that REEP2 is recruited to ERES, where it locates ERES on the Golgi and, in turn, promotes secretory trafficking. Interestingly, although ectopic REEP2 expression is sufficient to drive ERES/Golgi colocalization in epithelial cells (Fig. 4E vs. Fig. 4C, HCC827_Vec), REEP2 enhances the colocalization in a ZEB1-dependent manner (Fig. 4E). Whether this occurs via direct interactions between REEP2 and its interactors, which are co-opted by ZEB1, or indirectly through other altered membrane trafficking regulators in mesenchymal cells warrants to be determined.

Here, we identified a set of REEP2-dependent secreted proteins. ATX is aberrantly secreted by many human cancers to enhance cancer cell motility through lysophosphatidic acid receptors (LPARs) ^48–51^. Our recent study has shown that ATX is a key secretory effector within the EMT-driven secretome, contributing to cell motility ^11^. Furthermore, the ATX-CXCL1 axis recruits MDSCs while inhibiting cytotoxic CD8^+^ T-cell infiltration ^52^ and interleukin (IL)-6, a crucial regulator of MDSC accumulation and activation ^53^. REEP2 depletion suppresses the secretion of these factors. These findings are consistent with clinical data and our mouse model, however, it remains to be determined whether the REEP2-driven secretory program relates to LUAD progression and metastasis and how it correlates to EMT and the immune landscape of the TME in LUAD patients. In summary, our findings reveal REEP2 as a previously unrecognized effector of EMT-driven membrane trafficking program in LUAD, highlighting that REEP2-mediated ERES-Golgi coupling underlies the EMT-driven hypersecretory state.

## Methods

### Animal husbandry

For CRISPRi *in vivo* screen, all mouse studies were approved by the Institutional Animal Care and Use Committee at The University of Texas MD Anderson Cancer Center (Houston, Texas). Mice underwent standard care and were killed at predetermined time points or at first signs of morbidity according to the standards set forth by the Institutional Animal Care and Use Committee. To generate subcutaneous tumors, we injected 344SQ cells (1.1 × 10^6^ cells in 100 µL PBS) into the right flanks of syngeneic, immunocompetent 129-Elite (SOPF) mice (#476, Charles River Laboratories) (n = 6 per cohort). Tumor volumes were monitored weekly until tumors reached maximum size (∼2 cm^3^ total volume per mouse), and then the mice were sacrificed and the tumors were harvested, as previously described ^54^.

All other mouse studies were approved by the Institutional Animal Care and Use Committee at The University of Kentucky (Lexington, Kentucky). Mice underwent standard care and were killed at predetermined time points or at first signs of morbidity according to the standards set forth by the Institutional Animal Care and Use Committee. To generate orthotopic lung tumors, we intrathoracically injected 344SQ cells (0.5 × 10^6^ cells in 50 µL PBS) phosphate-buffered saline into the left lungs of syngeneic, immunocompetent 129-Elite (SOPF) mice (n = 10 per cohort) and carried out necropsies 14 d later to quantify primary tumor size and metastases to mediastinal lymph nodes and contralateral lung as we previously described ^11^. To generate subcutaneous tumors, we injected 344SQ cells (0.5 × 10^6^ cells in 100 µL PBS) into the right flanks of syngeneic, immunocompetent 129-Elite (SOPF) mice (n = 10 per cohort). Tumor volumes were measured weekly, and mice were sacrificed at 5 weeks post-injection, and necropsy was performed to quantify primary tumor weights and assess the presence of pleural surface lung metastases, as previously described ^11^.

### CRISPRi screens

The CRISPRi library virus preparation and screen were performed as previously described ^54^. In brief, a library of gRNAs targeting 2,099 known or putatively membrane trafficking genes with 5 gRNAs per gene for a total of 11,040 gRNAs (#83992, Addgene) ^27^ in the pCRISPRia-v2 vector (#84832, Addgene), and KRAB-dCas9-mCherry (#60954, Addgene), along with the psPax2 (Addgene cat# 12260) and pMD2.G (Addgene cat# 12259) lentiviral packaging vectors, were transfected into 293T cells for virus production by The Functional Genomics Core (The University of Texas MD Anderson Cancer Center, Houston, Texas). CRISPR library virus was used to titer on the murine mesenchymal LUAD 344SQ cells. The amount of virus that resulted in ∼30% cell positivity following cell sorting selection based on the BFP signal in the pCRISPRia-v2 vector was scaled up for library infection to ensure low multiplicity of infection (0.3) and appropriate library representation (500X for each sgRNA) after selection. Cells were infected in triplicate to generate biological replicates by The Functional Genomics Core.

On the day of animal injection, an initial time point (T0) of infected cells was harvested to determine a baseline of sgRNA abundance. For *in vivo* CRISPRi screens, three independent infections with the CRISPR library composed of biological replicates in the screening experiments. Each replicate was inoculated into the right flanks of 6 different 6-8-week-old 129-Elite (SOPF) mice with 1.1 million cells per injection. The tumors were harvested when they reached ∼2 cm^3^ total volume. The T0 infected cells and tumors were digested and the genomic DNA was extracted by The Biospecimen Extraction Facility (The University of Texas MD Anderson Cancer Center, Houston, Texas). The next-generation sequencing of sgRNA and analysis of sgRNA abundance were performed by VectorBuilder Inc. (Chicago, Illinois).

### Reagents

We purchased fetal bovine serum (FBS), live-cell imaging solution, phosphate-buffered saline (PBS), RPMI-1640, Trypsin-EDTA (0.25%), Alexa Fluor-tagged secondary antibodies, paraformaldehyde, bovine serum albumin (BSA), DAPI and Triton X-100, High-Capacity cDNA Reverse Transcription Kit, PowerUp™ SYBR™ Green Master Mix, and protease/phosphatase inhibitor cocktail from Thermo Fisher Scientific; Doxycycline (#D9891) from MilliporeSigma; Bronchial Epithelial Cell Growth Medium BulletKit from Lonza; jetPRIME transfection reagent from Polyplus Transfection; Transwell and Matrigel-coated Boyden chambers from BD Biosciences; Glass-bottom dishes and multiwell plates from MatTek; 10X Cell lysis buffer from Cell Signaling Technologies; 2X cell lysis buffer (#1610737) from Bio-Rad; CCK-8 (#K1018) from APExBIO; RNeasy Plus Mini Kit (#74136), and AllStars Negative Control siRNA (#1027281) from QIAGEN; Pre-miR™ miRNA Precursor Negative Control (miR-NC) (#AM17110); human miR-183a mimics (#PM12303) and human miR-193 mimics (#PM11786) from Thermo Fisher Scientific; human REEP2 siRNA #2 (#J-021201-18), and #4 (J-021201-20) from Horizon; primary antibodies against β-actin (#4967), ZEB1 (#3396) from Cell Signaling Technologies, REEP2 (#Ab191410) from Abcam, SEC16A (#20025-1-AP) from Proteintech, VSV-G (#EB0012) from Kerafast; Alexa Fluor 555 Mouse anti-GM130 (#560066) from BD Biosciences; REEP2-GFP (#RC202507L4) from Origene; SEC24D-mCherry (#32677), Golgi-pmTurquoise2 (#36205), mCherry-Golgi (#55052) and CD-MPR-GFP-RUSH (#202797) from Addgene; 3’-UTR luciferase reporters of REEP2 (#38786081) from Applied Biological Materials; pCI-Neo Renilla luciferase reporter (Basic) (#E1841) and pGL3 Basic Firefly luciferase control reporter (#E1751) from Promega; All-in-One miRNA qRT-PCR Detection Kit 2.0 (#QP115) from GeneCopoeia.

### Cell lines and culture

Murine LUAD cell lines were derived previously ^26^. Human LUAD cell lines were from Jonathan Kurie (The University of Texas MD Anderson Cancer Center, Houston, Texas). Human HBEC30KT cell line was from John Minna (The University of Texas Southwestern Medical Center, Dallas, Texas). Murine and human LUAD cell lines were cultured in RPMI-1640 with 10% FBS. Human HBEC30KT cell line was cultured in Bronchial Epithelial Cell Growth Medium BulletKit. Cells were maintained at 37°C in an incubator with a humidified atmosphere containing 5% CO2. Cells were transfected with vectors using jetPRIME transfection reagent.

### CRISPR/Cas9

CRISPR-Cas9-mediated murine REEP2 Knockout 344SQ cells were generated in the Cell-Based Assay Screening Service Core Facility (Baylor College of Medicine) using the following guide RNA sequences: 5′-aaagaugagccuagggaagc-3’ and 5′-aaacagaugggaguagagug-3’.

### Live and fixed cell imaging

Cells were seeded on type I collagen-coated cover glass (#1.5). Live cells were imaged in live-cell imaging media within incubation chambers. For fixed-cell imaging, cells were fixed using 4% paraformaldehyde for 10 min, permeabilized using 0.1% Triton X-100 for 5 min, and blocked with 5% BSA for 1h. Primary antibody incubation was performed in blocking buffer for 1 h overnight at 4°C, followed by Alexa Fluor-conjugated secondary antibodies in blocking buffer for 1 h at room temperature. Nuclei were counterstained with DAPI (#R37606, Thermo Fisher Scientific), and a cover glass was mounted using ProLong Diamond Antifade Mountant (#P36970, Thermo Fisher Scientific). Each step was followed by washing 3 times with PBS, which was also used as the solvent in all steps. For VSV-G assay, the cells were not permeabilized.

### VSV-G assay

The VSV-G transport assay was performed as described previously ^11^. In brief, cells were transiently transfected with EGFP-VSV-G (ts045), transferred to the restrictive temperature of 40°C for 20 h, and then transferred to the permissive temperature of 32°C for 1 h in the presence of 100 mg/ml cycloheximide, at which point the cells were fixed. In nonpermeabilized cells, exofacial and total VSV-G were detected by staining with an anti–VSV-G antibody and by measurement of EGFP signal intensity, respectively. VSV-G trafficking to the plasma membrane was measured based on the ratio of exofacial (surface) VSV-G fluorescence signal to the EGFP (total) signal intensity.

### RUSH assay

The RUSH assay was performed as described previously ^37,55^. In brief, cells were transiently transfected with a CD-MPR reporter, which is fused to the streptavidin-binding peptide (SBP), GFP-tag, and a second protein with a C-terminal ER retention signal (Lys-Asp-Glu-Leu; KDEL) fusing with streptavidin as a hook (#202797, Addgene). mCherry-Golgi (#55052, Addgene) was transiently co-transfected to label the Golgi compartment. BioLock (#2-0205-050, IBA Life Sciences) was used to quench excessive biotin in the culture medium according to manufacturer’s instructions. Twenty hours after transfection, the release of the RUSH cargos was induced by addition of 40 μM of D-biotin (#B4501, MilliporeSigma). Time-lapse acquisitions were done at 37°C in a thermostat-controlled Nikon stage top incubator (Nikon Instruments).

### Microscopy

Fixed and live-cell imaging were performed on Nikon CSU-W1 SoRa Confocal Microscope (Nikon Instruments) equipped with 60X/1.4 NA Oil, 100X/1.45 NA Oil objectives; 405/488/561/647 nm laser lines; GaAsP detectors, and Nikon stage top incubator. Images were acquired using NIS-Elements software (Nikon instruments).

### Image processing and quantitative analysis

For fixed cell imaging, the raw images were processed, and fluorescent intensity was analyzed in Fiji/ImageJ (www.imagej.nih.gov). For live cell imaging, the raw images were processed/analyzed in Imaris 10.0.1 (Bitplane software, Oxford instruments) with Spot features and MATLAB XTensions.

### Quantitative RT-PCR

Total RNA was isolated from cells using RNeasy Plus Mini Kit and subjected to reverse transcription and qPCR analysis as described ^56^. mRNA levels were normalized based on ribosomal protein L32 (RPL32) mRNA. MicroRNA levels were detected using the All-in-One miRNA qRT-PCR Detection Kit 2.0 with a universal reverse primer (#QP115, GeneCopoeia) according to the manufacturer’s instructions and normalized to U6. Primer sequences are listed in Table S3.

### Luciferase reporter assays

For 3′-UTR activity assays, H1299 cells were co-transfected with luciferase reporters driven by REEP2-3’-UTR sequences (100 ng) or pCI-Neo Renilla luciferase reporter (Basic) (100 ng), pGL3 Basic Firefly luciferase control reporter (100 ng), and microRNA mimics or control mimics (20 µM). After 48 hours, firefly and Renilla luciferase activities were measured with the Dual-Luciferase Reporter Assay System (Promega).

### Mutagenesis

Q5 Site-Directed Mutagenesis Kit (#E0554S, NEB) was used to generate mutants according to the manufacturer’s instructions. The primer sequences utilized for mutagenesis are: 5’-TCATTGGGTGGGGCTGAAAAAAACATGTTCCCACATTAAAA-3’ and 5’-CAGCCCCACCCAATGACCAGGAAAAGAG-3’ for human REEP2 3’UTR MT1 (miR-183 binding stie); 5′-AAAGAGGCCCCTCCTGAGGCTTCA-3′ and 5′-TTTAGCCCCTGCTCCACACTGTGC-3′ for human REEP2 3’UTR MT2 (miR-193a binding stie 1); 5′-AAATAGGGCCCAGTCTGCTTGGTACAG-3′ and 5′-TTTTCTGGCAGAACCCAGCCTCTG-3′ for human REEP2 3’UTR MT3 (miR-193a binding stie 2). All primers were purchased from Integrated DNA Technologies.

### Western blotting

Cells cultured in each well of a 6-well plate at 80% confluency were washed three times with PBS and harvested/resuspended in 150–200 μl of 2× Laemmli buffer (Bio-Rad). The cell lysate was boiled for 10 min and loaded onto an SDS gel. After transferring to a nitrocellulose membrane (Bio-Rad), membranes were blocked with 5% milk in TBST buffer and were probed with primary antibodies diluted in 5% BSA in TBST buffer. Horseradish peroxidase (HRP)-conjugated secondary antibodies were used according to the manufacturers’ instructions. Quantitative analysis was performed by using Fiji/ImageJ (NIH).

### Cell proliferation assay

Cells were seeded on 96-well plates at a density of 2*10^3^ cells per well on day 0. Relative cell density was measured by using the CCK-8 Counting Kit according to the manufacturer’s instructions. Briefly, cells were incubated with culture medium containing the CCK-8 solution for 2 h at 37 °C, and the absorbance was read at 450 nm.

### Cell cycle assay

Cells were synchronized using culture medium containing 0.1% FBS for 24 h, then allowed to grow in culture medium containing 10% FBS for 24 h. The cells were fixed with 70% ethanol at 4°C overnight, and were incubated with Propidium Iodide (PI)/RNase Staining Solution (#4087S, Cell Signaling Technologies) for 30 min. After incubation, cell cycle was analyzed using a flow cytometer (BD Symphony).

### Cell apoptosis assay

Cells were harvested and then incubated with 7-AAD (#420404, BioLegend) and Annexin-V (#640926, BioLegend) at room temperature for 10-20 min. After incubation, cell apoptosis was analyzed using a flow cytometer (BD Symphony).

### Cell migration and invasion assays in Boyden chambers

As described previously ^11^, 1*10^5^ cells were seeded in the upper wells of Transwell for migration and Matrigel-coated Boyden chambers for invasion assays, respectively (BD Biosciences) and allowed to migrate toward 10% FBS in the bottom wells. After 16 h of incubation, migrating or invading cells were stained with 0.1% crystal violet, photographed, and counted. For conditioned medium (CM) transfer experiments, CM samples were isolated, filtered through a 0.45-μm filter, and applied to cells seeded in the upper wells.

### Liquid chromatography/mass spectrometry (LC-MS)

The immunoprecipitation (IP) from agarose beads was then washed with ice-cold lysis buffer 5 times. The bound proteins on beads were eluted in 5% SDS. For mass spectrometry analysis, samples were reduced, alkylated, and digested using trypsin on an S-trap as described ^57^. The resultant peptides were quantified and normalized within sets by the Quantitative Fluorometric Peptide Assay. Samples were analyzed using a 90-minute gradient on our Orbitrap Eclipse. The resulting Liquid chromatography-mass spectrometry (LC-MS) data was processed in Proteome Discoverer 2.5 and SEQUEST using a human database. The minora node was used to perform Label-Free Quantitation (LFQ) using the MS peak areas for each of the peptide-spectral matches. The IP-MS was performed in triplicate.

### Immune cell profiling

Mouse flank tumors generated by subcutaneous injection of 0.5*10^6^ parental 344SQ or 344SQ_REEP2_KO #25 or #44 cells into the flank of wild-type mice for 5 weeks were harvested. The tumor tissues and mouse spleens were processed for isolating tumor-infiltrating lymphocytes by The Flow Cytometry and Immune Monitoring Core Facility (University of Kentucky, Lexington, Kentucky). Single-cell suspension was stained as described previously ^11^ with the following antibodies: CD3-PE-594 (#100246, BioLegend), CD45-Pacific Blue (#103126, BioLegend), CD4-APCCy7 (#100526, BioLegend), CD8-PE-Cy7 (#100721, BioLegend), CD278-PE (#117406, BioLegend), CD25-BUV395 (#564022, BD Biosciences), PD1-BV-505 (#135220, BioLegend), CD62L-FITC (#35-0621-U500, Thermo Fisher Scientific), CD44-BV-711 (#103057, BioLegend), TIM3-APC (#134007, BioLegend), FoxP3-PerCP-Cy5.5 (#45-5773-82, Invitrogen), GR1-BV-711 (#106443, BioLegend), CD11b-BV-650 (#101239, BioLegend), CD11c-BV-785 (#117335, BioLegend), F4/80-APC (#204801-U100, Tonbo), PDL1-PE-Dazzle-594 (#124323, BioLegend), CD86-APC-Cy7 (#105030, BioLegend), MCHII-PE-Cy7 (#107629, BioLegend), CD80-BV-605 (#104729, BioLegend), CD68-PerCP-Cy5.5 (#137009, BioLegend), iNOS-PE (#125920-80, Invitrogen), Arg1-FITC (IC5868F, R&D Systems), CD31-BV-786 (#740870, BD Biosciences), and the live/dead cell marker Ghost Violet-BV-510 (#10-0870-T100, VWR). For intracellular staining, cells were fixed and permeabilized using the intracellular staining perm wash buffer (BioLegend) according to manufacturer instructions. Data were acquired using a flow cytometer (BD Symphony) and analyzed using FlowJo software (version 10.10.0; BD Biosciences).

### Analysis of Kaplan-Meier Survival Data

LUAD patient survival data were downloaded from the Kaplan-Meier plotter database ^58^. Analysis of LUAD patients was performed in REEP2 high- and low-expression cohorts. The *P* value was calculated using the log-rank test ^58^.

### TNMplot analysis

The REEP2 mRNA expression analysis in normal lung and different stages of lung tumor tissues was performed as previously described ^59^.

### Statistical analysis

Unless mentioned otherwise, the results shown are representative of replicated experiments and are the means ± standard error of the means (SEM) from triplicate samples or randomly chosen cells within a field. The comparison of mRNA levels with EMT scores and the correlation analysis in human lung cancers were performed as previously described ^60^, and the EMT score was calculated as previously described ^60,61^. Statistical evaluations were carried out with Prism 9 (GraphPad Software, Inc.). Unpaired 2-tailed Student t-tests or ANOVA were used to compare means for two or more groups, respectively, and *P* values < 0.05 were considered statistically significant.

## Data, materials, and software availability

All data are available in the main text or the supporting information.

## Acknowledgement

This work was supported by the National Institutes of Health (NIH) through R00 CA249048 (to G.X.), R03 CA280382 (to X.T.), R01 CA181184 (to J.M.K.), R01 CA2111125 (to J.M.K.), NIH Lung Cancer SPORE grant P50 CA70907 (to J.M.K.), and T32CA165990 (to K.F.). We thank The Functional Genomics Core (The University of Texas MD Anderson Cancer Center) for CRISPRi library virus preparation and cell infection, and The Biospecimen Extraction Facility (The University of Texas MD Anderson Cancer Center) for genomic DNA extraction from CRISPRi in vivo screen. This research was supported by the Flow Cytometry and Immune Monitoring (FCIM) Shared Resource of the University of Kentucky Markey Cancer Center (P30CA177558). We thank the FCIM for tumor tissue processing for immune cell profiling.

## Author contributions

K.F., O.O., and G.X. conceived, designed, executed, and interpreted all experiments. S.W. and X.T. assisted with mutagenesis for 3’-UTR reporter assays. X.L. and J.Y. assisted with mouse experiments for CRISPRi *in vivo* screen. J.M.K. supported and supervised the CRISPRi *in vivo* screen. J.X. assisted with CRISPR/Cas9-mediated genome-editing cell line establishment. W.K.R. directed and interpreted mass spectrometry experiments. X.T. and G.X. supervised the project and contributed to the design and interpretation of all experiments.

## Figures

**Fig. S1.**
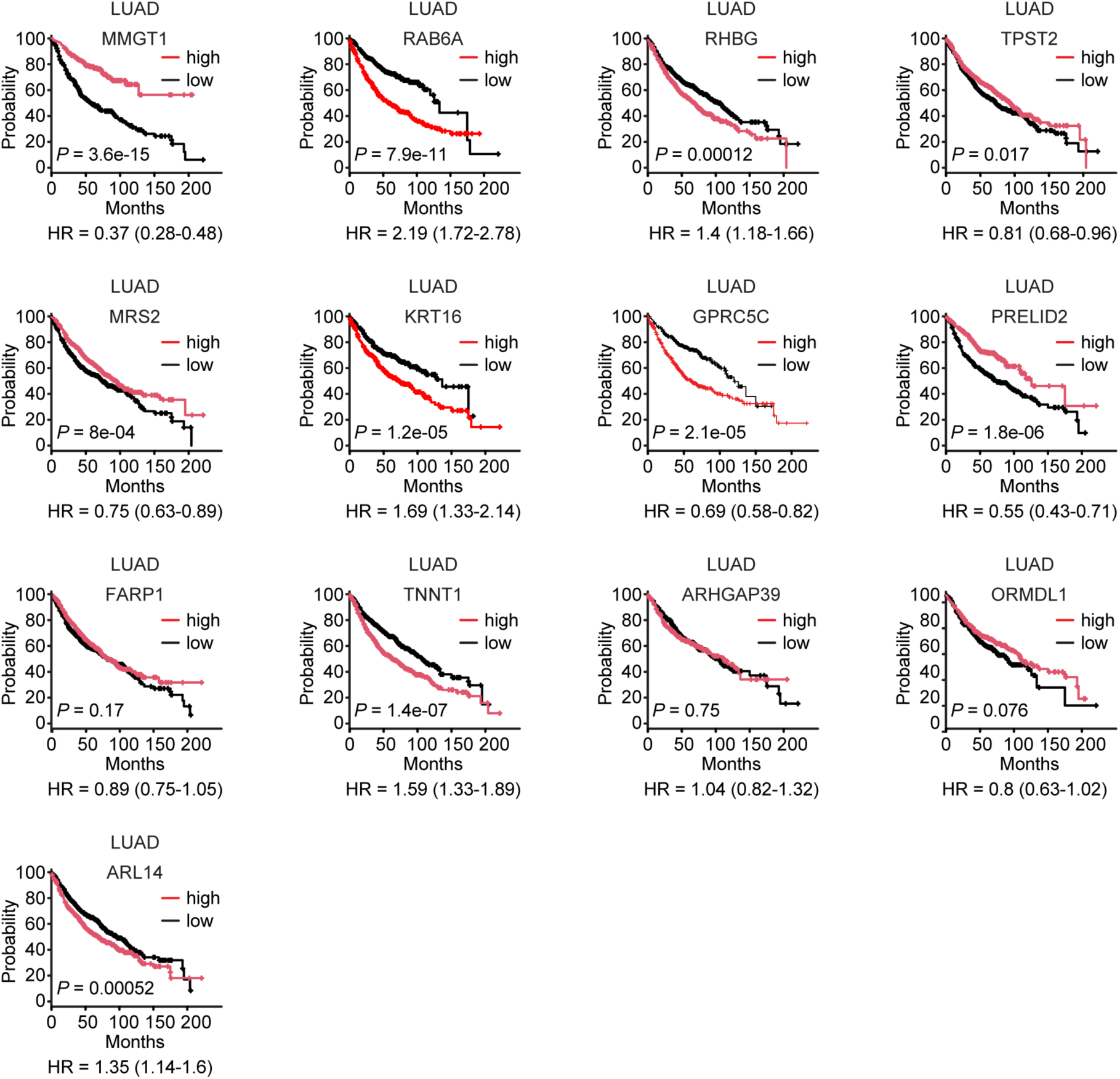
Kaplan-Meier survival analysis of LUAD patients based on the mRNA levels of indicated genes above (high) or below (low) the median value.

**Figure S2.**
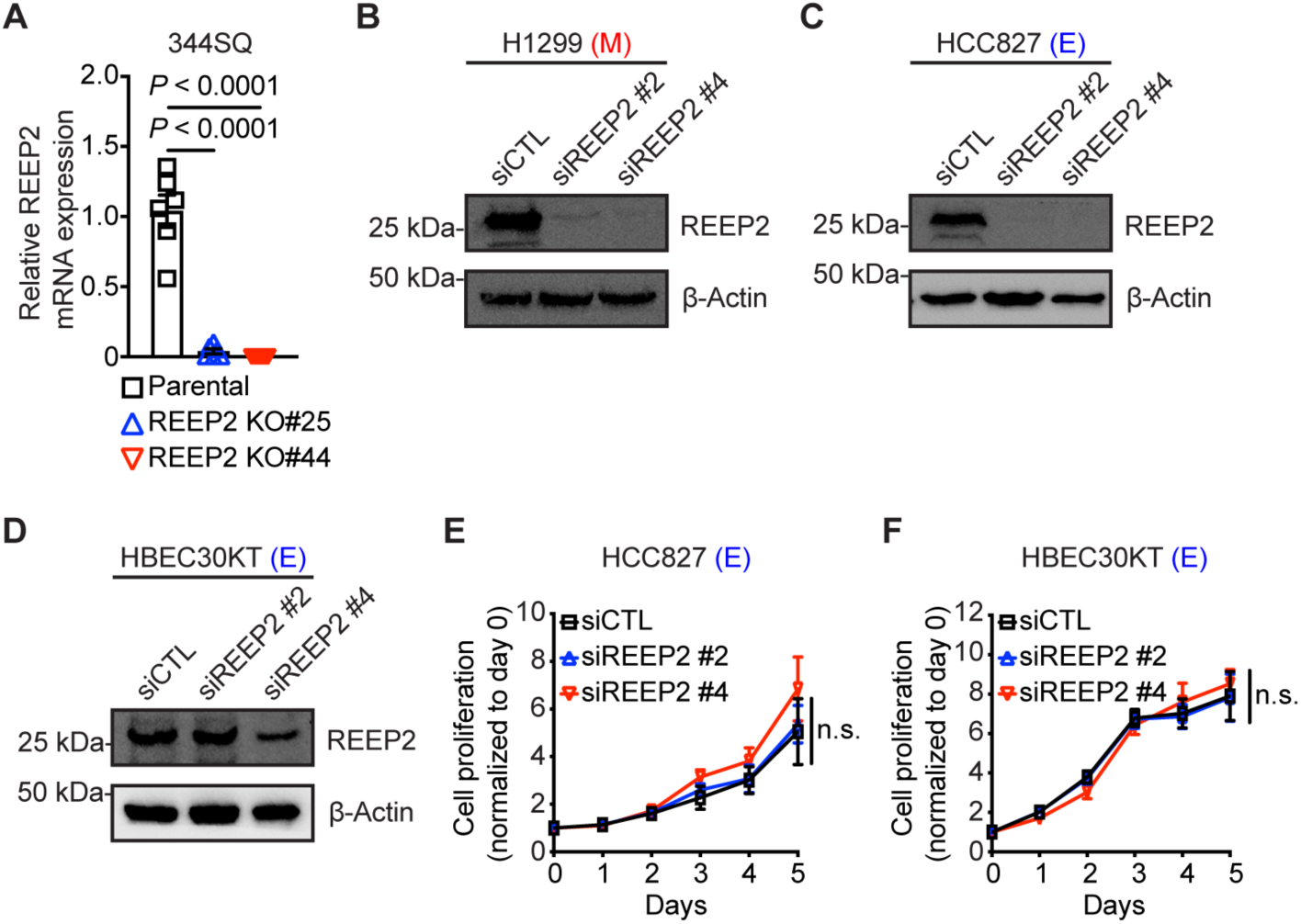
Depletion of REEP2 in murine and human LUAD cells. (A) qPCR analysis of REEP2 mRNA expression levels in parental or REEP2 knockout (KO) 344SQ cell lines (n=6 replicates per condition). (B-D) WB analysis of REEP2 levels in human H1299 (B), HCC827 (C), and HBEC30KT (D) cells transfected with control siRNA (siCTL) or REEP2 siRNA (siREEP2). β-Actin loading control. M: mesenchymal. E: epithelial. (E, F) *In vitro* cell proliferation assay in human HCC827 (E) and HBEC30KT (F) cells transfected with control siRNA (siCTL) or REEP2 siRNA (siREEP2). Results represent means ± SEM. *P* values were determined using two-tailed Student’s t-test.

**Figure S3.**
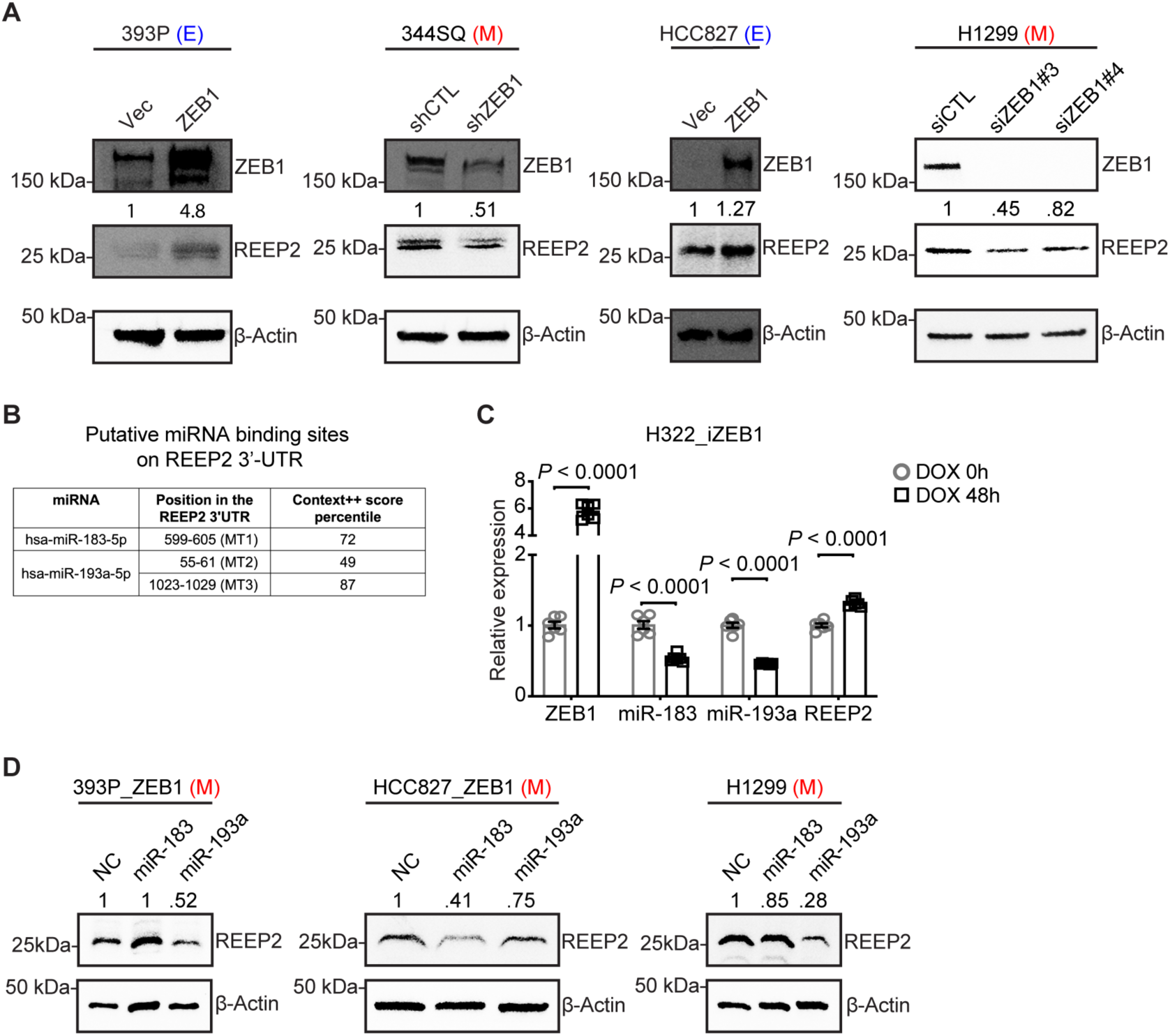
ZEB1 upregulates REEP2 through the regulation of miRNAs. (A) WB analysis of ZEB1 and REEP2 protein expression levels in murine (393P, 344SQ) and human (HCC827, H1299) LUAD cells with ectopic ZEB1-expression or ZEB1-depletion. β-actin loading control. Relative densitometric values are indicated. E: epithelial. M: mesenchymal. (B) Putative miRNA binding sites on REEP2 3’UTR predicted by TargetScan and used for mutagenesis (MT1-3) (C) qPCR analysis of ZEB1, miR-183, miR-193a, and REEP2 expression levels in human H322_iZEB1 cells with doxycycline (DOX) (1 µg/ml) treatment (n=6 replicates per condition). Results represent means ± SEM. *P* values were determined using two-tailed Student’s t-test. (D) WB analysis of REEP2 protein expression levels in the indicated cells transfected with miR-183 or miR-193a mimics or non-coding control (NC). M: mesenchymal.

**Figure S4.**
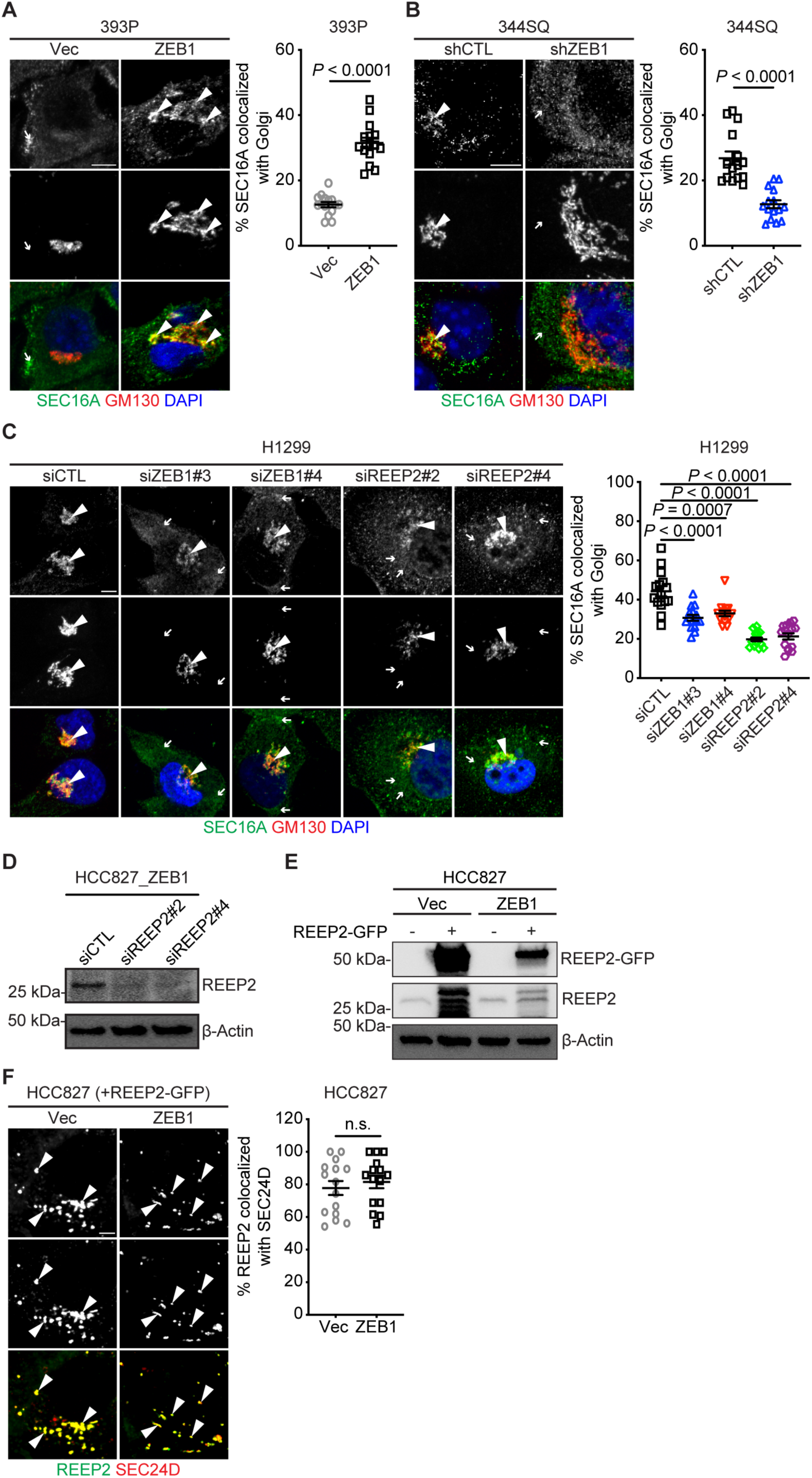
ZEB1 drives ERES colocalization with the Golgi via REEP2. (A-C) Confocal micrographs of cells co-stained with anti-SEC16A (green) and anti-GM130 (red) antibodies. DAPI (blue). Scale bar: 5 μm. The scatter plots quantify Golgi-localized SEC16A per cell (dot) based on % of total SEC16A that co-localizes with Golgi (GM130 channel) (n = 15 cells per group) in murine epithelial cells (393P) with ectopic ZEB1-expression (A), in murine mesenchymal cells (344SQ) with ZEB1 depletion (B), or in human mesenchymal cells (H1299) with ZEB1 or REEP2 depletions (C). (D) WB analysis of REEP2 protein expression levels in HCC827_ZEB1 cells transfected with control siRNA (siCTL) or REEP2 siRNA (siREEP2). β-actin loading control. (E) WB analysis of REEP2 protein expression levels in HCC827 cells co-transfected with empty vector (Vec) or ZEB1, with REEP2-GFP. β-actin loading control. (F) Confocal micrographs of cells co-transfected with REEP2-GFP (green) and SEC24D-mCherry (red). Scale bar: 5 μm. The scatter plots quantify SEC24D-localized REEP2 per cell (dot) based on % of total REEP2 that co-localizes with SEC24D (n = 15 cells per group) in HCC827 cells with ectopic ZEB1-expression. Results represent means ± SEM. *P* values were determined using two-tailed Student’s t-test.

**Figure S5.**
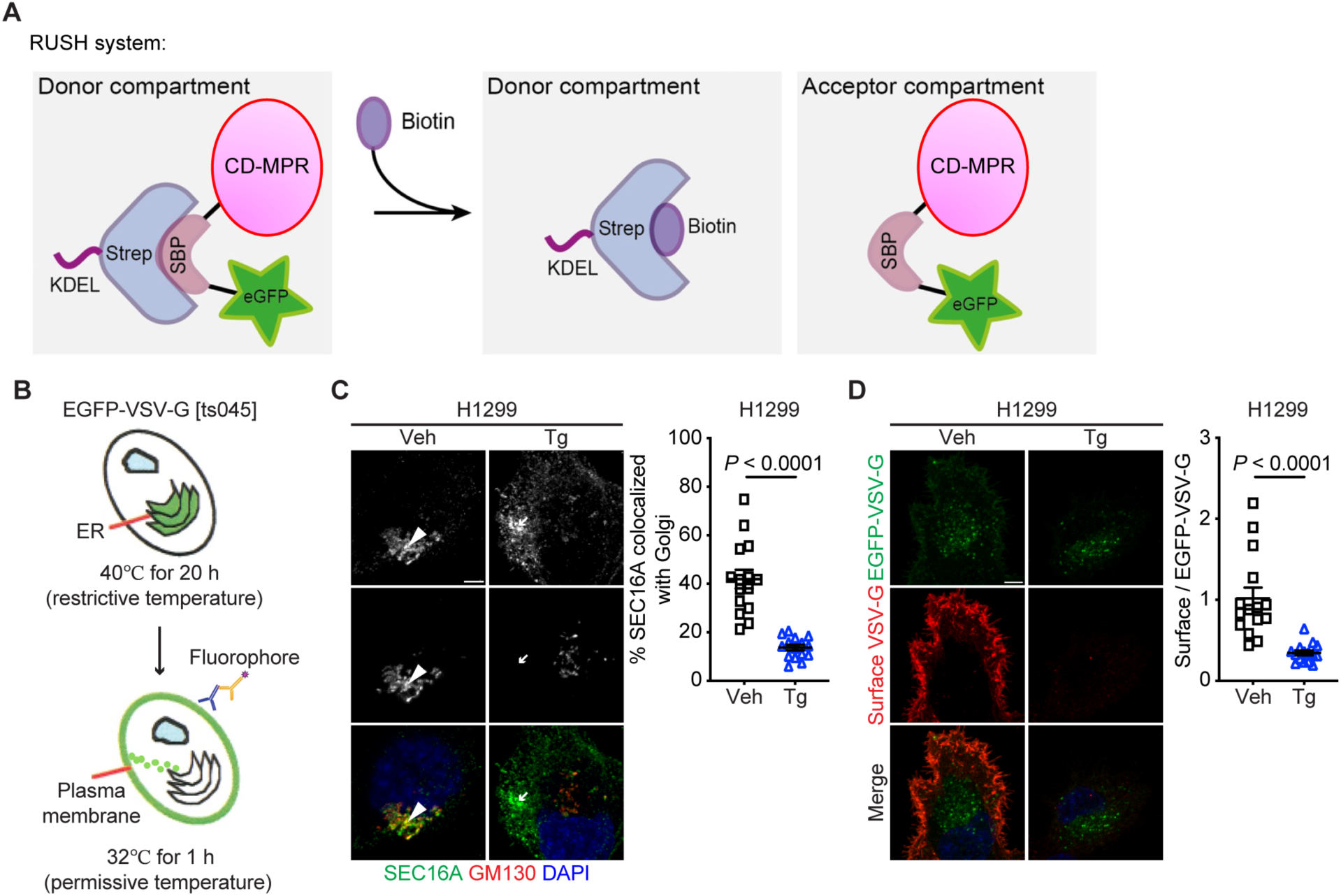
ERES/Golgi colocalization promotes secretory trafficking. (A) Schematic illustration of the retention using selective hooks (RUSH) system. The cells were co-transfected with the reporter protein (CD-MPR), which is fused to the streptavidin-binding peptide (SBP) and GFP, and a second protein with a C-terminal ER retention signal (Lys-Asp-Glu-Leu; KDEL) fusing with streptavidin as a hook. The interaction of CD-MPR with the hook protein retains CD-MPR in the ER compartment and biotin administration releases the CD-MPR-GFP reporter from the ER and transfers it to the Golgi. (B) Schematic of VSV-G assay. The cells were transfected with vectors that express an EGFP-tagged temperature-sensitive mutant VSV-G (EGFP-VSV-G [ts045]) that is transported from the ER to the plasma membrane via the Golgi. At designated time points after switching cells to temperatures that cause VSV-G accumulation (40°C) and release (32°C) from the ER, cells were fixed and exofacial VSV-G was detected in non-permeabilized cells by staining with anti-VSV-G monoclonal antibody. The VSV-G trafficking to the plasma membrane is determined by the ratio of exofacial (surface) VSV-G fluorescence signal to the EGFP signal intensity. (C) Confocal micrographs of cells co-stained with anti-SEC16A (green) and anti-GM130 (red) antibodies. DAPI (blue). Scale bar: 5 μm. The scatter plots quantify Golgi-localized SEC16A per cell (dot) based on % of total SEC16A that co-localizes with Golgi (GM130 channel) (n = 15 cells per group) in H1299 cells treated with vehicle control (Veh) or 5 µM thapsigargin (Tg) for 1 h. (D) Confocal micrographs of EGFP-VSV-G-transfected cells taken 1 h after transfer to permissive temperature. Scale bar, 5 μm. The scatter plot represents the ratio of surface VSV-G to EGFP-VSV-G in each cell (dot) (n = 15 cells per group) in H1299 cells treated with vehicle control (Veh) or 5 µM thapsigargin (Tg) for 1 h. Results represent means ± SEM. *P* values were determined using two-tailed Student’s t-test.

**Figure S6.**
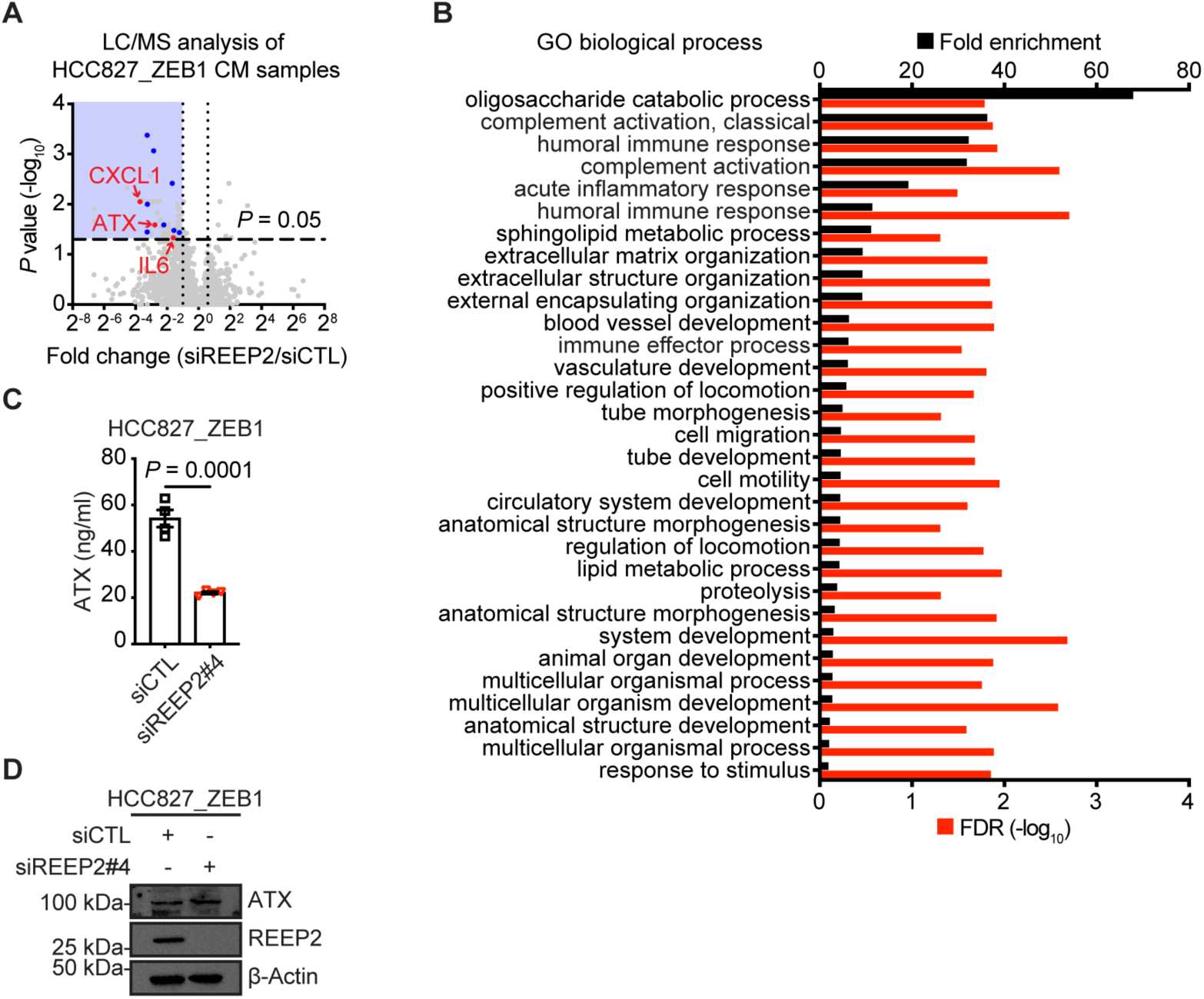
REEP2 activates a pro-metastatic secretion. (A) Volcano plot of proteins identified by LC/MS analysis of CM samples. *P* values (Y axis) and fold-change (X axis). Blue box indicates proteins with significantly reduced secretion (blue box, *P* < 0.05) in REEP2-depleted HCC827_ZEB1 cells (siREEP2). The blue and red dots label the factors that are associated with immune regulation. (B) Enrichment analysis (Fisher’s exact test using Gene Ontology terms) of the down-regulated secreted proteins. (C) ELISA assay shows the ATX levels in CM samples (n = 4 replicates per group). *P* values were determined using two-tailed Student’s t-test. (D) WB analysis of ATX and REEP2 levels in human HCC827_ZEB1 cells transfected with control siRNA (siCTL) or REEP2 siRNA (siREEP2#4). β-actin loading control.

